# Outbreak of Highly Pathogenic Avian Influenza Virus H5N1 in Seals in the St. Lawrence Estuary, Quebec, Canada

**DOI:** 10.1101/2023.11.16.567398

**Authors:** Stéphane Lair, Louise Quesnel, Yohannes Berhane, Pauline Delnatte, Carissa Embury-Hyatt, Marie-Soleil Nadeau, Oliver Lung, Shannon T. Ferrell, Robert Michaud

**Affiliations:** Université de Montréal, St. Hyacinthe, Quebec, Canada; Canadian Food Inspection Agency, Winnipeg, Manitoba, Canada; Ministère de l’Agriculture, des Pêcheries et de l’Alimentation du Quebec, St. Hyacinthe, Quebec, Canada; Groupe de recherche et d’éducation sur les mammifères marins, Tadoussac, Quebec, Canada

## Abstract

We describe an unusual mortality event caused by a highly pathogenic avian influenza virus (HPAI) H5N1 clade 2.3.4.4b involving harbor (*Phoca vitulina*) and grey (*Halichoerus grypus*) seals in the St. Lawrence Estuary, Quebec, Canada. Fifteen (56%) of the seals submitted for necropsy were considered to be fatally infected by H5N1 containing fully Eurasian (EA) or Eurasian/North American genome constellation.

Concurrently, presence of large numbers of bird carcasses infected with H5N1 at haul-out sites most likely contributed to the spill-over of infection to the seals. Histologic changes included meningoencephalitis (100%), fibrinosuppurative alveolitis, and multi-organ acute necrotizing inflammation. This is the first report of fatal H5N1 infection in pinnipeds in Canada, raising concerns about the expanding host of this virus, potential for establishment of a marine mammal reservoir, and the public health risks associated with spillover to mammals.

---

Sporadic outbreaks of influenza A virus (IAV; genus *Alphainfluenzavirus*) infections have been reported in pinnipeds in the United States and Europe, most commonly in harbor (*Phoca vitulina*) and grey (*Halichoerus grypus*) seals *(1–3)*. The characterization of the IAV subtypes associated with these outbreaks has repeatedly suggested an avian-variant origin thought to occur through a cross-species transmission, or spillover between wild aquatic birds acting as reservoirs for IAV and susceptible mammalian hosts *(4, 5)*. So far, the reported outbreaks of IAV in pinnipeds have been mainly associated with fatal respiratory diseases *(6, 7)*. Harbor seals seem to be particularly susceptible to IAV infections, and factors such as close contact with wild birds and specific mammalian adaptations of the virus subtypes have been suggested as drivers in establishing a potential reservoir of IAV in marine mammal populations *(5, 8)*.

Although epidemics of IAV have been reported since the late 1970s in seals on the North American Atlantic coast, fatal infections by Alphainfluenzavirus have never been documented in marine mammals from the St. Lawrence Estuary and Gulf, Quebec, Canada. Moreover, despite the documented high seroprevalence of influenzavirus A and B (genus *Betainfluenzavirus*) in Canadian seal populations *(9)*, seal mortality caused by an influenza infection had never been reported in Canadian waters until recently, when a novel low pathogenicity avian IAV (H10N7) caused a fatal bronchointerstitial pneumonia in a harbor seal in British Columbia, Canada *(10)*.

In December 2021, A/goose/Guangdong/1/1996 (Gs/GD) lineage HPAI H5N1 (2.3.4.4b clade) was detected in Eastern Canada in wild gulls *(11)*. This novel virus subtype was likely introduced by wild birds that carried it across the Atlantic Ocean through pelagic routes or during direct winter migration *(11)*. In the following months, this 2.3.4.4b clade H5N1 virus spread rapidly across North America, reassorted with North American lineage IAVs causing unprecedented outbreaks in many species of wild birds and commercial and backyard poultry flocks, with spillover to several species of wild terrestrial mammals *(12–16)*.

During the summer of 2022, deaths of harbor and grey seals caused by 2.3.4.4b clade H5N1 were confirmed in Eastern Quebec, Canada and on the coast of Maine, USA, prompting the National Oceanic and Atmospheric Administration (NOAA) to declare an Unusual Mortality Event (UME) for Maine harbor and grey seals *(17)*. HPAI infections have also recently been reported as the cause of mortality of harbor seals in the North Sea (H5N8) *(18)*, grey seals in the Baltic Sea (H5N8) *(19)*, and South American sea lions (*Otaria flavescens*) *(20)* in Peru. However, this is the first report of infections and mortality by HPAI A(H5N1) in marine mammals in Canada and the first detailed description of H5N1-induced lesions in pinnipeds. Given the emergence of this 2.4.4.4b clade H5N1 virus in marine mammal species of the St. Lawrence Estuary and the public health concerns associated with mammalian spillover of avian IAV, we aim to describe the 2022 outbreak of HPAI A(H5N1) affecting pinniped species, with an emphasis on epidemiological data and pathology findings.

## Material and Methods

### Stranding Data Analysis

Stranding data were obtained via the archives of the Quebec Marine Mammal Emergency Response Network. The network monitors mortality and morbidity of marine mammals in the St. Lawrence River, Estuary and Gulf (48°23’N 69°07’W). The number of stranded (mortality or morbidity) harbor seals, grey seals, and seals of unspecified species during the second and third trimesters of 2022 (from April 1 to September 30) was compared to the average of the 10 previous years for the same period using a one-sample t-test. Goodness-of-fit of the stranding distribution of 2012 to 2021 was evaluated with a Shapiro-Wilk test for both groups of seals. Distribution was considered normal if P > 0.05. Seals of unidentified species were combined with harbor seals for this comparison to control for the differences in species identification rates in 2022 compared to previous years (data not shown).

### Postmortem Examination

Selection of carcasses to be examined was based on the state of decomposition *(21)* and field access. All carcasses, which were all submitted frozen, were examined after thawing by veterinary pathologists with experience in marine mammal pathology. Animals were classified in 2 age groups, < 1 year-old or adult, based on their total length (22). Animals with no evidence of muscular or fat depletion were considered to be in good nutritional condition. Tissue samples of major organs, including lung, heart, kidney, brain, intestines, lymph nodes, liver, spleen, pancreas, tongue, adrenal gland, esophagus, bladder, stomach, thyroid gland, mammary gland and thymus, were processed for histopathologic evaluation by light microscopy using standard laboratory procedures. For each necropsy case, a separate nasal swab and a rectal swab were collected using a sterile polyester-tipped plastic applicator (UltiDent Scientific, https://www.ultident.com) placed in UTM™ medium (Micronostyx, https://micronostyx.com). Additionally, rectal and nasal swabs were collected from seals found stranded for which the carcass could not be examined either due to poor preservation state or logistical limitations. All samples were first tested for IAV by PCR at the provincial Animal Health Laboratory. All IAV H5-positive samples were subsequently sent to the National Centre for Foreign Animal Disease (NCFAD) laboratory in Winnipeg for confirmatory testing. Samples of brain or lung were also submitted for PCR for cases with lesions suggestive of IAV, but that tested negative on swabs.

### RNA Extraction, RT-PCR, and Virus Isolation

Both laboratories (MAPAQ and NCFAD) used the same methods for RNA extraction and PCR. Total RNA was extracted from clinical specimens (swabs and tissues) and virus isolates using the MagMax AM1836 96 Viral RNA Isolation Kit (ThermoFisher Scientific, https://www.thermofisher.com) as per the manufacturer’s recommendation using the KingFisher Duo prime, KingFisher Flex, or Apex platforms (ThermoFisher Scientific). The spiked Enteroviral armoured RNA (ARM-ENTERO; Asuragen, https://asuragen.com) was used as an exogenous extraction and reaction control. The extracted RNA samples were tested for the presence of influenza A virus genomic material using the matrix gene-specific real-time reverse transcription PCR (rRT-PCR) assay. Samples positive with the matrix primer set were subsequently tested with the H5- and H7-specific rRT-PCR primer sets as described previously (23, 24). Tests with a Ct value below 36.00 were considered positive, and tests with a Ct value between 36.00 and 40.00 were considered suspicious. For virus isolation, PCR-positive samples were inoculated into 9-day-old embryonated specific pathogen-free (SPF) chicken eggs via the allantoic route.

### Nanopore Sequencing and Genome Assembly

To determine the clade, lineage, and clusters of each positive sample, the full genome segments of IAVs were amplified directly from clinical specimens or isolates using RT–PCR as described previously *(25)*. Nanopore sequencing was performed on an Oxford Nanopore GridION sequencer with an R9.4.1 flowcell after library construction using the rapid barcoding kit (SQKRBK004 or SQK-RBK110.96). The raw Nanopore signal data was basecalled and demultiplexed with Guppy (v5.1.12) using the high accuracy or super-accurate basecalling model on each runs. Basecalled Nanopore reads were analysed and assembled with the CFIANCFAD/BLAST (v2.12) search of IRMA assembled genome segment sequences against all sequences from the NCBI Influenza Virus Sequence Database (n = 959,847) (https://www.ncbi.nlm.nih.gov/genomes/FLU/Database/nph-select.cgi?go) and influenza virus sequences from the 2021/2022 H5N1 outbreaks.

### Immunohistochemistry

For immunohistochemistry (IHC), paraffin tissue sections were quenched for 10 minutes in aqueous 3% hydrogen peroxide. Epitopes were then retrieved using proteinase K for 15 minutes and then rinsed. The primary antibody applied to the sections was a mouse monoclonal antibody specific for influenza A nucleoprotein (NP) (F26NP9, produced in-house) and was used at a 1:5,000 dilution for 30 minutes. The primary antibody binding was then visualized using a horse radish peroxidase labelled polymer, Envision® + system (anti-mouse) (Dako, https://www.agilent.com/en/dako-products) and reacted with the chromogen diaminobenzidine (DAB). The sections were then counter stained with Gill’s haematoxylin.

## Results

### Stranding Data

From April 1 to September 30, 2022, a total of 209 dead or sick seals were reported in the waters bordering the province of Quebec, Canada: 127 harbor seals, 47 grey seals, 6 harp seals (*Pagophilus groenlandicus*), 1 hooded seal (*Cystophora cristata*), and 28 seals of undetermined species. The number of dead or sick stranded harbor seals and seals of unknown species combined (155) was 3.7 higher than the average number for these 2 groups combined in the previous 10 years for the same interval (N=41.6), representing a statistically significant increase in number of strandings (t = 5.55, Df = 9, P-value < 0.001) (Figure 1A). A statistically significant increase of similar magnitude was noted for the number of grey seals found dead or sick during the second and third trimesters of 2022 (47 in 2022 compared to an average of 12 in 2012-2021; t = 4.35, Df = 9, P-value = 0.002) (Figure 1B). No increase in mortality or morbidity were observed for the other species of pinnipeds compared to the previous 10 years.

**Figure 1.**
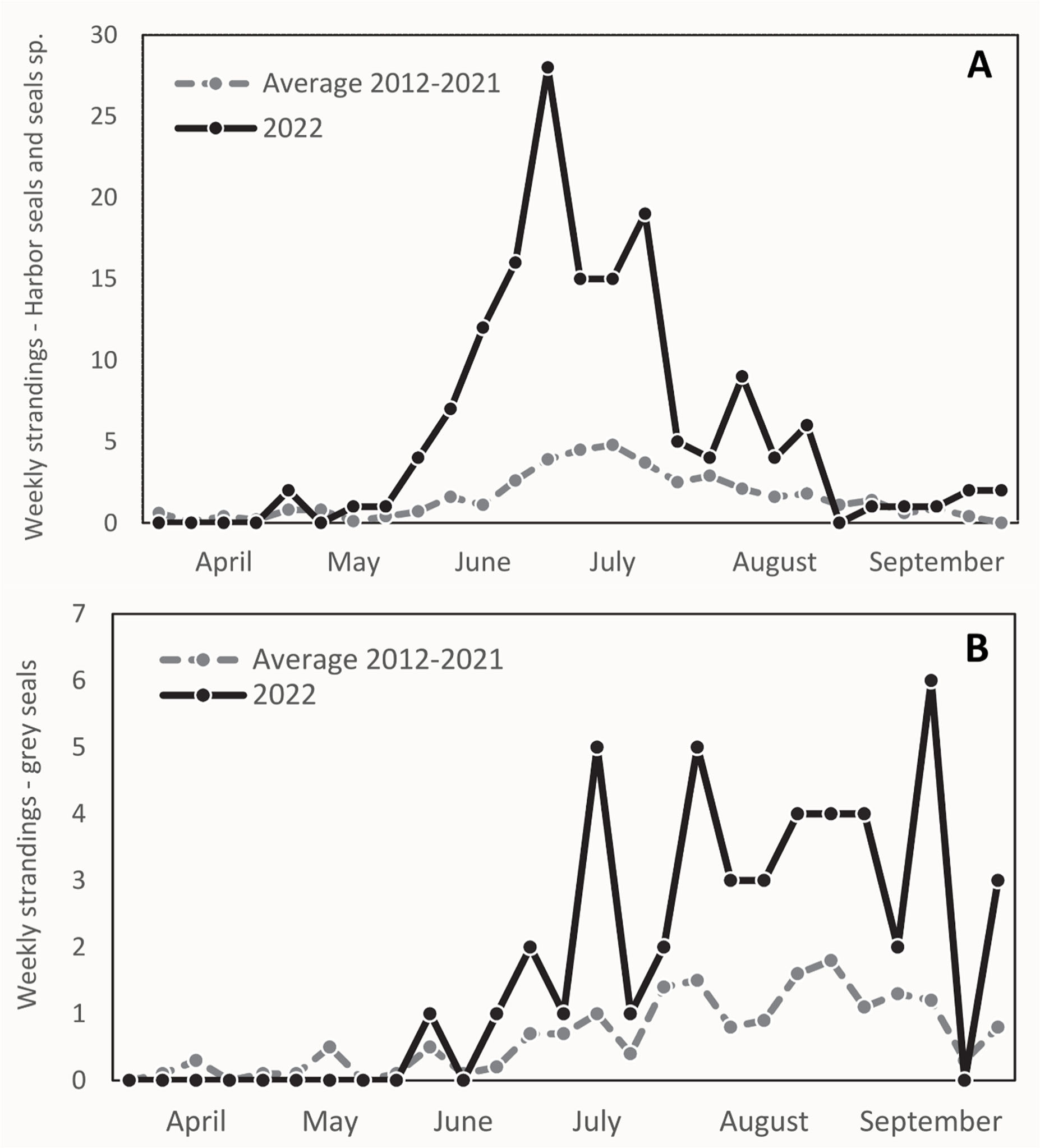
Weekly occurrences of stranded seals from April 1, 2022, to September 30, 2022, in the St. Lawrence Estuary and Gulf, Quebec, Canada, compared to the average number of strandings over the previous 10 years (2012 to 2021) during the same trimesters. A) Harbor seals and seals of undetermined species. B) Grey seals.

### Descriptive Epidemiology and Necropsy Findings

Carcasses of 22 harbor seals, 3 grey seals, 1 harp seal, and 1 hooded seal found between April 22 and October 6, 2022 were submitted for postmortem examination. Based on molecular detection of H5 and presence of lesions suggestive of IAV infection, HPAI A(H5N1) was identified to have caused the death of 14 harbor seals and 1 grey seal (56%) (Table 1). Combined nasal and rectal swabs were sampled on the shore from an additional 11 harbor seals and 1 grey seal carcasses. HPAI A(H5N1) was identified in 6 of the harbor seals (50%) (Table 1).

**Table 1.** Percentage of stranded seals tested between April 22^nd^ and October 6^th^, 2022 that were infected by HPAI A(H5N1) in the St. Lawrence Estuary, Quebec, Canada.

| Species | Percentage of cases of infection by HPAI A(H5N1)*<br>(number infected†/number tested) |
| --- | --- |
| **Harbor seal (*Phoca vitulina*)** |  |
| Full postmortem examination | 64% (14/22) |
| Nasal/rectal swab (field) | 55% (6/11) |
| Total | 61% (20/33) |
| **Grey seal (*Halichoerus grypus*)** |  |
| Full postmortem examination | 33% (1/3) |
| Nasal/rectal swab (field) | 0% (0/1) |
| Total | 25% (1/4) |
| **Hooded seal (*Cystophora cristata*)** |  |
| Full postmortem examination | 0% (0/1) |
| **Harp seal (*Pagophilus groenlandicus*)** |  |
| Full postmortem examination | 0% (0/1) |
| **Total pinnipeds** |  |
| Full postmortem examination | 56% (15/27) |
| Nasal/rectal swab (field) | 50% (6/12) |
| Total | 54% (21/39) |

\*HPAI A(H5N1): Highly pathogenic avian Influenza virus type A subtype H5N1.
†Cases of HPAI A(H5N1) infections were determined based on epidemiological data, presence of suggestive lesions, and PCR testing (using H5 specific primers).

All 21 infected seals were found between May 30 and July 8, 2022 in the estuarine segment of the St. Lawrence waterway, mainly on the south shore, between the towns of Baie-Comeau (49.2213° N, 68.1504° W) and Notre-Dame-du-Portage (47.7630° N, 69.6096° W) (Figure 2). Infections were detected in both < 1 year-old and adult seals with no age predilection. All the infected adult harbor seals (n=9) were female and 6 of them had evidence of recent parturition (either active lactation and/or asymmetrical uterine horns without the presence of a fetus). The infected adult grey seal was a male. There were 3 males and 7 females (and 1 non-determined) infected <1 year-old seals. Age classes and sex of the infected seals are presented in Table 2. One of the infected seals was found alive with profound lethargy and neurological signs. In addition, anecdotal observations of weak and dyspneic harbor seals were reported during the outbreak.

**Figure 2.**
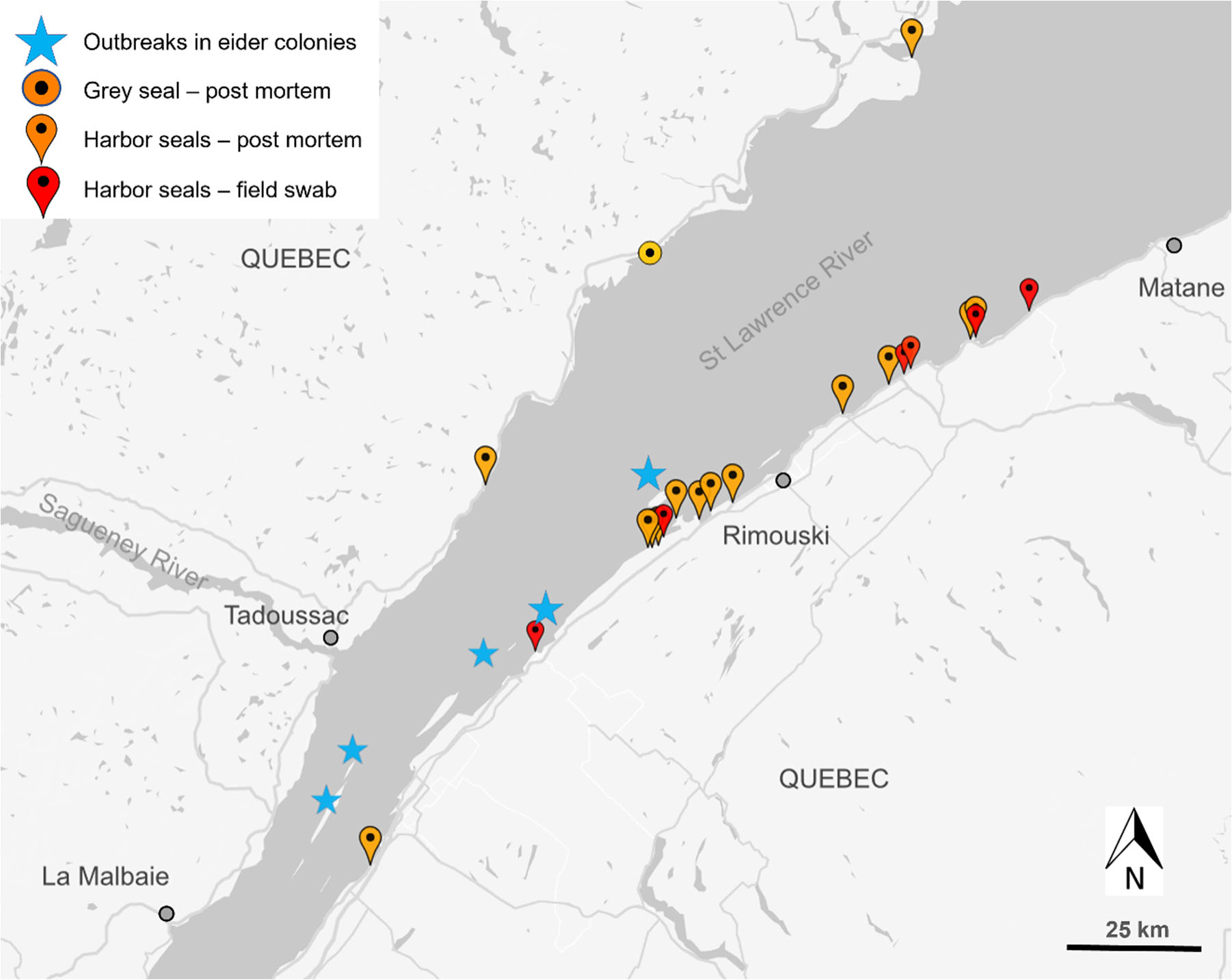
Geographical locations of stranded dead or sick seals infected by HPAI A(H5N1) during the 2022 outbreak in the St. Lawrence Estuary, Quebec, Canada. The locations of the documented outbreaks in common eider colonies are also presented.

**Table 2.**
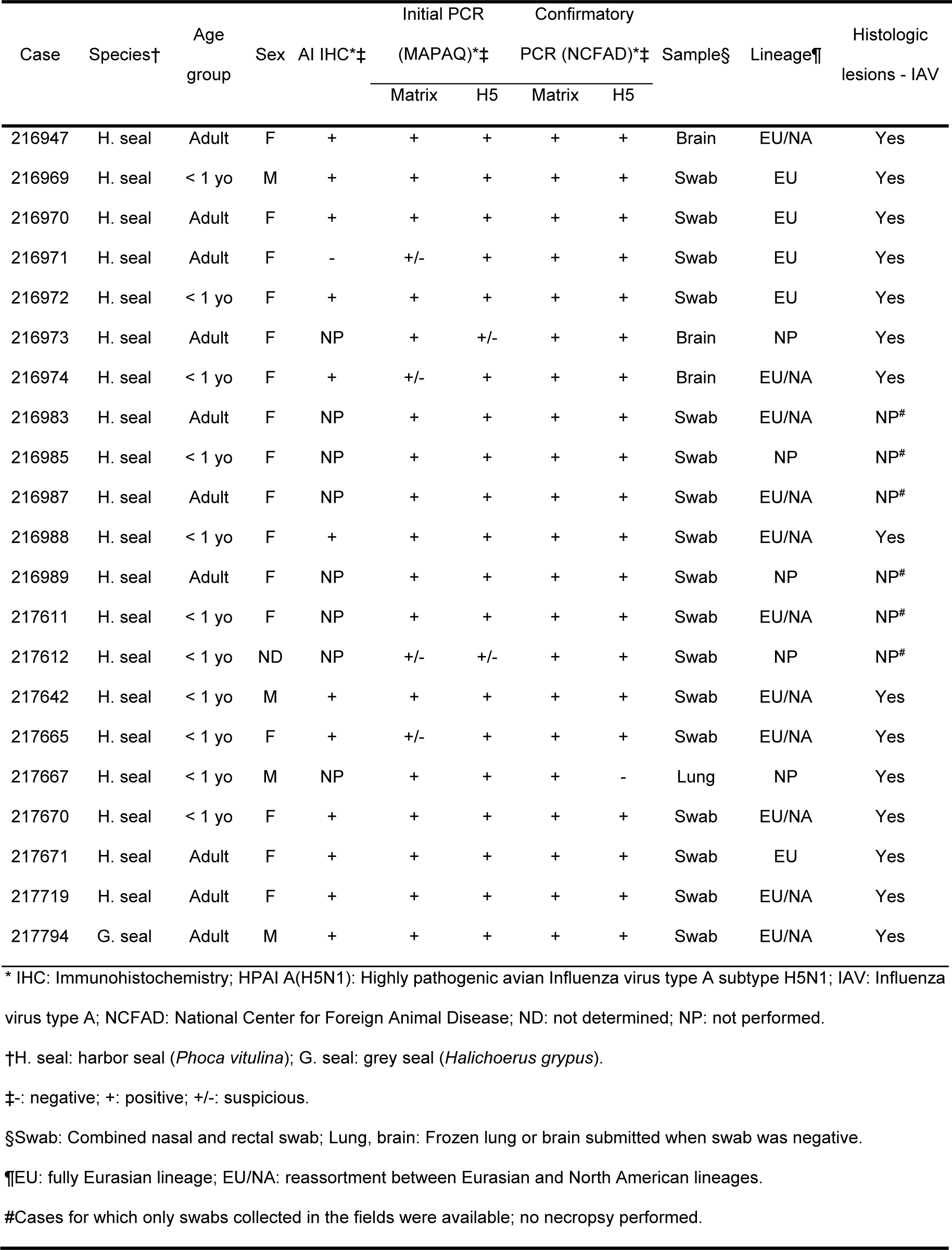
Demographic data and results of immunohistochemistry and molecular testing performed in seals infected by HPAI A(H5N1) during the 2022 outbreak in the St. Lawrence Estuary, Quebec, Canada.

Carcasses were attributed decomposition scores *(21)* varying from code 2 – 2.5 (fresh to mild decomposition) in 12 cases and to code 3 (moderate decomposition) for 3 cases. Twelve seals were in excellent nutritional condition, 2 were thin and 1 was emaciated. The main postmortem findings are reported in Table 3. Notable gross lesions were limited to lymphadenomegaly (submandibular and mesenteric lymph nodes), red-tinged foam in the tracheal lumen, and pulmonary congestion. The occurrences of relevant histologic lesions are presented in Table 3. Acute multifocal to diffuse mixed meningoencephalitis (Figure 3A), characterized by a predominantly neutrophilic infiltrate with lymphocytes in Virchow-Robin spaces, the meninges, and/or the submeningeal neuropil, was present in all cases. Neuronal necrosis, satellitosis, and gliosis were also regularly observed. Acute fibrinosuppurative alveolitis, consisting of neutrophils with fibrinous aggregates in the alveolar lumen, was observed in 9/15 seals (Figure 3B). Interstitial pneumonia, characterized by a mild to moderate mixed inflammatory infiltrate within the alveolar septae, was seen either concurrently or distinctively from the alveolitis. In addition to the inflammatory changes, alveolar emphysema, mild acute alveolar damage with hyaline membranes, and necrotic type-1 pneumocytes were sometimes present. Six animals presented multifocal necrotic foci of the adrenal cortex with infiltrates of degenerate neutrophils (Figure 3C). Acute necrotizing thymitis, lymphadenitis, and splenitis were also observed and consisted of multifocal to coalescing zones of necrosis, mainly centered on cortical lymphoid follicles along with extensive lymphoid depletion (Figure 3D). Lymph nodes were often reactive with an accumulation of histiocytic cells in the lymphatic trabeculae and medullary sinus, macrophages in the germinal centers, and an infiltration of the subcapsular sinus with granulocytes and macrophages. Other microscopic changes included 6 cases with mild membranous glomerulonephritis, 4 cases with necrotizing fibrinous hepatitis of variable severity, 1 case of mild multifocal myocarditis with degenerate myofibers, and 1 animal with mild neutrophilic perivascular infiltration in the perimysium.

**Figure 3.**
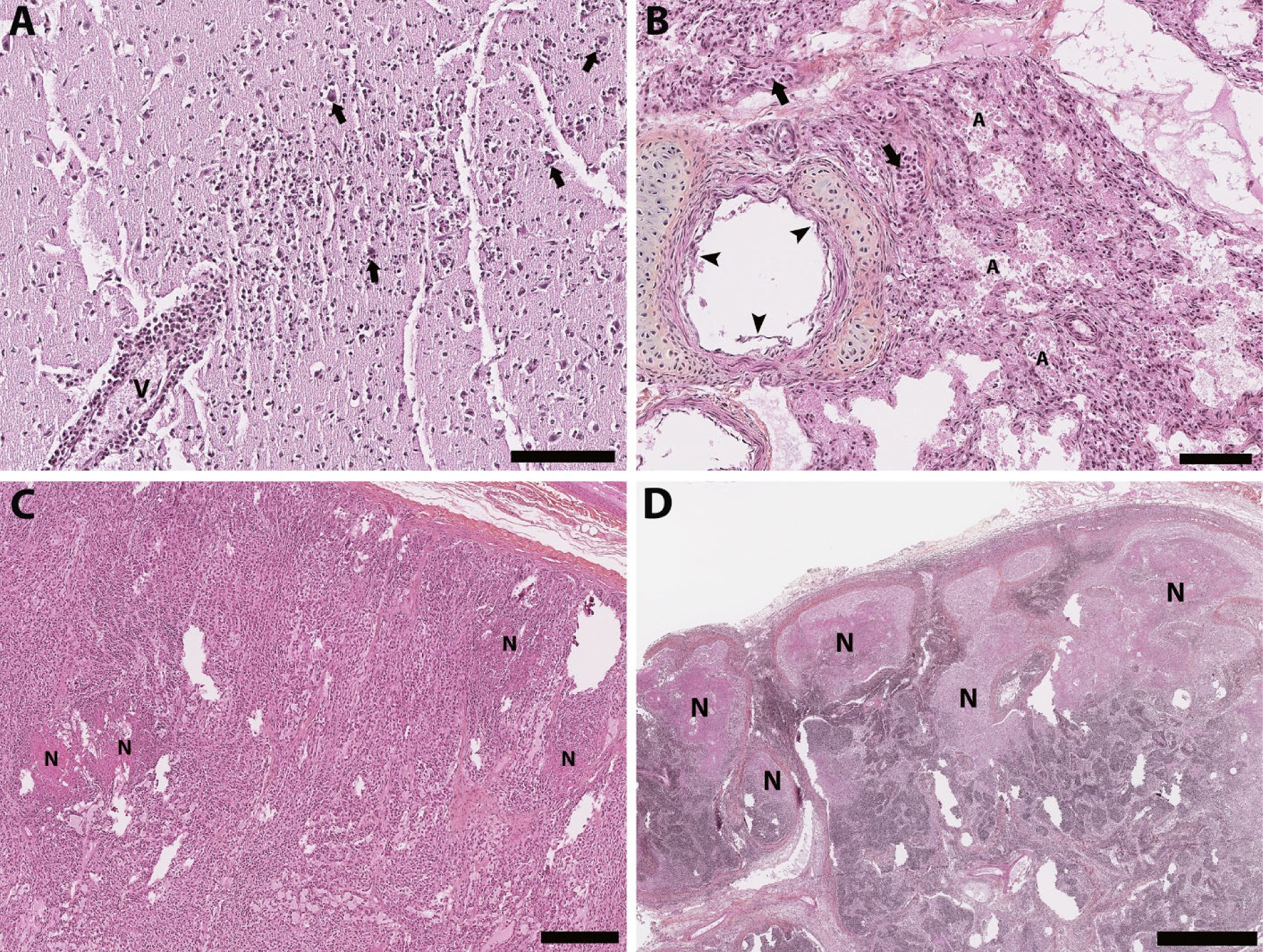
Histology section of thawed, formalin-fixed tissues from harbor seals (*Phoca vitulina*) infected by HPAI A(H5N1). Hematoxylin phloxine saffron stain. A) Brain of a young of the year female. The Virchow-Robin space around a vessel (V) is infiltrated by numerous layers of polymorphonuclear cells. Several neurons have a condensed hyperacidophilic cytoplasm indicative of necrosis and are often associated with satellitosis (arrows). A focally extensive infiltration of the neuropil by neutrophils and glial cells is also present. Bar = 100 µm. B) Lung from a young of the year female. The alveolar (A) and vascular (arrow) lumens contain numerous often degenerate polymorphonuclear cells. The alveolar walls are infiltrated by numerous inflammatory cells composed of neutrophils and mononuclear cells. The epithelial cells bordering the small bronchi are often necrotic. Bar = 100 µm. C) Adrenal gland of an adult female. Multifocal foci of necrosis are present in the cortical zone (N). Bar = 300 µm. D) Lymph node from an adult female. Marked multifocal to coalescing necrosis of the lymphoid tissues in the cortical region (N). Bar = 1 mm.

**Table 3.** Microscopic lesions documented in harbor (*Phoca vitulina*) and grey seals (*Halichoerus grypus*) infected by HPAI A(H5N1) during the 2022 outbreak in the St. Lawrence Estuary, Quebec, Canada.

| Microscopic lesions | Percentage of affected seals<br>(number with lesions/number examined) |
| --- | --- |
| Brain |  |
| Meningoencephalitis | 100% (15/15) |
| Predominantly neutrophilic | 80% (12/15) |
| Predominantly lymphoplasmacytic | 20% (3/15) |
| Lung |  |
| Pulmonary inflammatory changes | 73% (11/15) |
| Acute multifocal fibrinosuppurative alveolitis | 60% (9/15) |
| Acute interstitial pneumonia | 53% (8/15) |
| Acute alveolar damage | 40% (6/15) |
| Alveolar emphysema | 27% (4/15) |
| Adrenal gland |  |
| Adrenocortical necrosis | 60% (6/10) |
| Thymus |  |
| Acute multifocal necrotizing thymitis | 50% (2/4) |
| Lymph nodes |  |
| Reactional lymph nodes | 43% (6/14) |
| Acute multifocal necrotizing lymphadenitis | 36% (5/14) |
| Kidney |  |
| Multifocal membranous glomerulonephritis | 40% (6/15) |
| Liver |  |
| Necrotizing hepatitis | 29% (4/14) |
| Spleen |  |
| Necrotizing splenitis | 21% (3/14) |
| Muscle |  |
| Neutrophilic infiltration of the perimysium | 9% (1/11) |
| Heart |  |
| Multifocal myocarditis | 7% (1/14) |

Immunoreactivity for influenza A virus antigen was observed in tissues from 12 of the 13 cases that were tested (Table 2). Variable abundance of antigens, from mild to extensive, were detected in different tissues, including the neuropil and neurons (Figure 4A), pulmonary alveolar septa and glandular bronchial cells (Figure 4B), renal glomeruli, spleen, pancreas, liver, vascular walls of skeletal muscle, lymph nodes, tracheal vessels, and the adrenal gland. These antigens were often associated with, but not limited to, necrotic foci.

**Figure 4.**
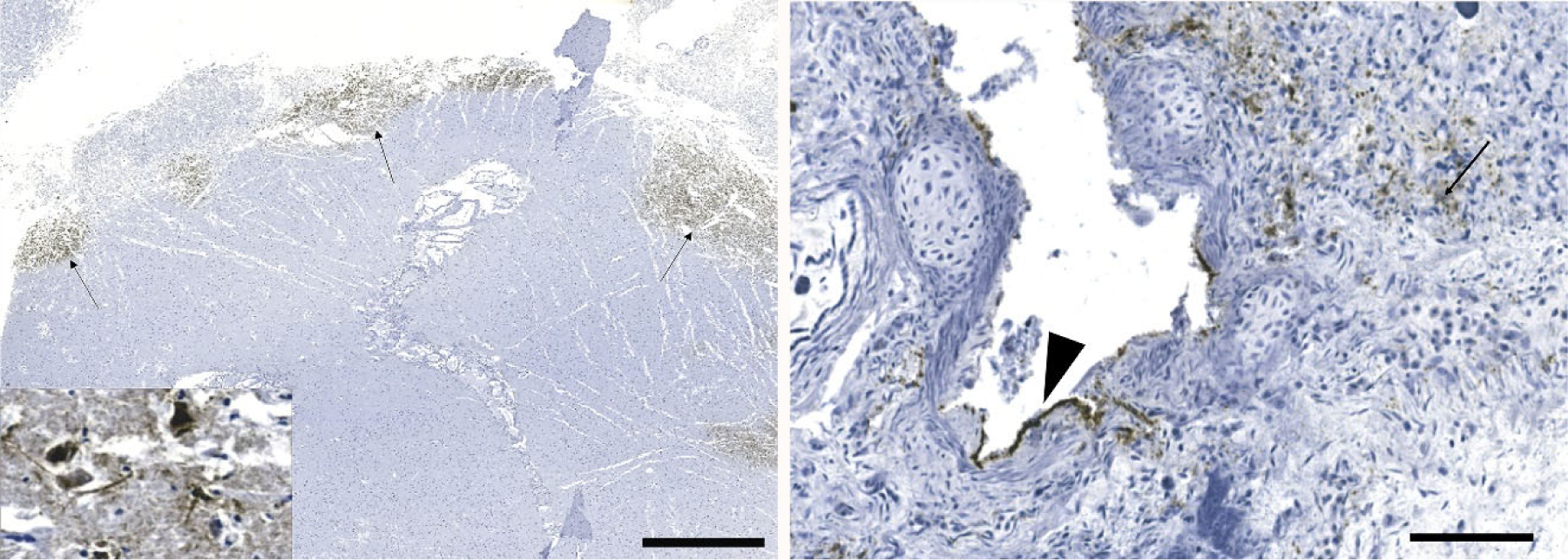
Detection of influenza virus antigen by immunohistochemistry in brain (A) and lung (B) of harbor seals (*Phoca vitulina*) infected by HPAI A(H5N1). A) There are multifocal areas of intense immunostaining (arrows) with staining of all structures in the affected area including neurons and neuropil (inset). Bar = 2 mm. B) Positive immunostaining can be observed within alveolar septae (arrow) and in bronchiolar epithelial cells (arrowhead). Bar = 80 µm.

### Virology Assessment

The presence of H5 RNA was confirmed by PCR in 15 seals on which a necropsy was performed and in 6 seals that were swabbed in the field (n=21). All samples were negative for H7. All the 15 necropsied seals had lesions suggestive of IAV on histology. In 4 cases, NCFAD confirmatory PCRs for H5 performed on combined rectal/nasal swabs were negative but H5 RNA was detected by subsequent PCR on frozen lung or brain tissues. Virus isolation (using embryonated SPF chicken eggs) was successful in 16 of those 21 cases. The 16 isolates were sequenced to determine the subtype and lineage of the H5 virus. All isolated belonged to Gs/GD lineage H5N1 viruses belonging to the to the 2.3.4.4b clade. Five isolates had fully Eurasian gene segments similar to the Newfoundland (NL)-like 2.3.4.4b clade H5N1 viruses that emerged in Canada in late 2021. The other 11 isolates were reassortment H5N1 viruses containing gene segments PB2, PB1, PA and NP belonging to the North American lineage IAVs and gene segments HA, NA, M and NS belonging to the NL-like 2.3.4.4b clade H5N1 viruses. Detailed results of the molecular testing are presented in Table 2.

## Discussion

Investigation of the increased mortalities in the pinniped populations of the St. Lawrence Estuary and Gulf during the summer of 2022 detect H5N1 infections for the first time in harbor and grey seals in Canadian waters. All necropsied seals that were positive for H5 by PCR also presented histologic lesions consistent with IAV infection. The demonstration by IHC that influenza A virus antigens were often associated with necrotic and inflammatory lesions further supports a causal relationship between this virus, the lesions observed, and the death of these animals. It is reasonable to assume that the almost 4-fold increase in summer mortalities in harbor and grey seals compared to historic data could be explained, at least in part, by a H5N1 outbreak affecting these populations of seals.

Even if good agreement was observed between the different diagnostic modalities, there were a few discrepancies between the results that are worth noting (Table 2): IHC failed to detect one infected seal; 4 cases that were considered only suspect based on an initial matrix or H5 PCRs (MAPAQ) were later confirmed positive subsequently (NCFAD); and in 4 cases, combined nasal/rectal swabs tested negative on PCR while frozen tissues (lung or brain) tested positive. These differences in diagnostic results highlight the importance of a multi-step diagnostic methodology to confirm avian IAV cases.

Most of the examined cases were in good nutritional condition with only minor macroscopic changes, except for noticeable lymphadenomegaly. However, it is possible that some macroscopic changes were overlooked due to postmortem and freezing artefacts. Inflammatory and necrotic changes were detected in several organs, including brain, lungs, adrenal glands, liver, and lymphoid organs. The central nervous system (100%) and the lungs (73%) were the most commonly affected organs. This neurotropism and respirotropism are similar to what has been reported in birds *(26)* and in mesocarnivores infected with H5N1 *(12, 15)*, as well as in three harbor seals from the German North Sea infected with HPAI H5N8 *(18)*. The predominantly neutrophilic nature of the meningoencephalitis is unusual for a viral infection and indicative of a very acute infection.

The route of transmission and the viral pathogenesis for HPAI infection have not been established in pinnipeds. In most mammals, including humans, IAVs are transmitted via inhalation of aerosols or respiratory fomites from an infected individual and replicate primarily in respiratory epithelium. Previous reports of infections with low pathogenicity avian influenza (LPAI) in pinnipeds are consistent with the same pathophysiology as with stranded sick seals displayed respiratory signs, while dead seals showed necrotizing hemorrhagic bronchitis and alveolitis *(6, 7)*. However, the clinical presentation and the postmortem lesions seen in various mammalian species affected by H5N1, including the seals from this study, indicate a clear neurotropism *(12, 15, 16)*. Neurological signs, encephalitis, and a high load of viral antigens and RNA in the brain all point towards a different pathogenesis and possibly, a different route of inoculation. To date, most mammalian species that have been reported infected by H5N1 are carnivores that are likely to prey or scavenge on wild birds. Consequently, ingestion of birds infected with HPAI is presumed to be the most likely source of infection in free-ranging wild mesocarnivores *(12, 15)* and in sporadic cases of infected domestic or captive wild carnivores *(27, 28)*. This route of transmission, which has been proven successful with red foxes being experimentally infected with HPAI H5N1 clade 2.2 after eating infected bird carcasses *(29)*, is also plausible in grey seals since this species is known to prey on sea birds, making a weakened bird or infected carcass a potential source of infection. However, this pathway of infection is unlikely for harbor seals since they are not known to predate or scavenge on birds. Consequently, transmission via environmental exposure, such as accidental ingestion of feces or feathers from infected birds, drinking of fecal-contaminated water, or inhalation of aerosols or respiratory fomites from infected avian carcasses should be considered for harbor seals. All harbor seals that died of HPAI A(H5N1) in this population were found in relatively close proximity to islands used as pupping ground by this population. Some of these islands also harbor breeding colonies of marine birds, such as the common eiders (*Somateria mollissima*). Numerous cases of mortality caused by HPAI A(H5N1) were documented in these colonies of marine birds during the same time period (Canadian Wildlife Health Cooperative database). Anecdotal observations of seals hauling out on sites where carcasses of eiders were present were reported during the outbreak. These observations would then support the hypothesis that harbor seals were exposed to HPAI A(H5N1) following close contact with infected marine birds. The fact that the infected harbor seals were either newborn or females in breeding age would also supports the link between these infections and the contact with marine birds at pupping sites. The presence of two 2.3.4.4b H5N1 viruses with two different genome constellations of HPAI and the temporo-geographical distribution of the seals would indicates that this outbreak was associated with more than one source of infection. With the data currently available, it is not possible to determine if direct seal-to-seal transmission occurred.

The infection of mammalian species, such as seals, by HPAI A(H5N1) viruses raises concern about recent viral mutations facilitating the entry and replication within mammalian cells. From a human health perspective, such changes in viral host range warrant continued vigilance to detect a potentially deadly epidemic before its emergence. In addition, marine mammals, such as seals or other pinnipeds, might act as reservoirs for this virus, which could contribute in increasing the risk of mutations and viral reassortment, favoring the infection of new mammalian hosts. It is, therefore, essential to monitor the occurrence and molecular characteristics of this HPAI virus in populations of wild marine mammals in order to better understand the public health risk associated with this emerging pathogen – host dynamic.

## Acknowledgements

The authors would like to thank the staff and volunteers of the RQUMM and the staff of CWHC-QUEBEC for their help with carcasses retrieval and necropsy. Thanks also to the professional and technical staff of the CDEVQ for their help with sample analysis, and to Estella Moffat for the technical assistance with immunohistochemistry. Finally, the authors acknowledge Tristan Juette for his assistance with statistics, and Jean-Francois Giroux and Matthieu Beaumont for their contribution regarding the unusual mortality event in common eiders.

We are grateful for the funding provided to O. Lung by the Canadian Safety and Security Program (CSSP-2018-TA-2362) for the Oxford Nanopore GridION sequencer. The scientific conclusions and opinions expressed herin are those of the authors and do not necessarily reflect the views or policies of the Canadian Government, its agencies, or any of the included organizations.

